# Evolutionary emergence of infectious diseases in heterogeneous host populations

**DOI:** 10.1101/317099

**Authors:** Hélène Chabas, Sébastien Lion, Antoine Nicot, Sean Meaden, Stineke van Houte, Sylvain Moineau, Lindi M. Wahl, Edze R. Westra, Sylvain Gandon

**Author notes:** Corresponding author: CEFE UMR 5175, CNRS - Université de Montpellier - Université Paul-Valéry Montpellier – EPHE, 1919 route de Mende, 34293 Montpellier, France.

## Abstract

Emergence and re-emergence of pathogens are notoriously difficult to predict. The erratic nature of those events is reinforced by the stochastic nature of pathogen evolution during the early phase of an epidemic. For instance, mutations allowing pathogens to escape host resistance may boost pathogen spread and promote emergence. Yet, the ecological factors that govern such evolutionary emergence remain elusive both because of the lack of ecological realism of current theoretical frameworks and the difficulty of experimentally testing their predictions. Here we develop a theoretical model to explore the effects of the heterogeneity of the host population on the probability of pathogen emergence, with or without pathogen evolution. We show that evolutionary emergence and the spread of escape mutations in the pathogen population is more likely to occur when the host population contains an intermediate proportion of resistant hosts. We also show that lower pathogen inoculum size and higher diversity of host resistance decrease the probability of evolutionary emergence. Crucially, we present experimental confirmations of these predictions using lytic bacteriophages infecting their bacterial hosts containing diverse CRISPR-Cas immune defenses. We discuss the implications of these results for cross-species spillover and for the management of emerging infectious diseases.

**Significance statement:** Can we predict the emergence of infectious diseases? The probability that an epidemic breaks out is highly dependent on the ability of the pathogen to acquire new adaptive mutations and to induce evolutionary emergence. Forecasting pathogen emergence thus requires a good understanding of the interplay between epidemiology and evolution taking place at the onset of an outbreak. Here, we provide a comprehensive theoretical framework to analyze the impact of host population heterogeneity on the probability of pathogen evolutionary emergence. We use this model to predict the impact of the fraction of susceptible hosts, the inoculum size of the pathogen and the diversity of host resistance on pathogen emergence. Our experiments using lytic bacteriophages and CRISPR-resistant bacteria support our theoretical predictions.

## Introduction

The emergence and re-emergence of infectious diseases is a major public-health concern. Understanding the factors that govern the ability of pathogens to invade a new host population is of paramount importance to design better surveillance systems and control policies. Mathematical epidemiology can provide key insights into these dynamics (1–4). For instance, simple deterministic models identified critical vaccination thresholds above which pathogens are driven extinct, which informed policy guidelines for vaccination campaigns (1). However, chance events and rapid pathogen evolution can also play a critical role in determining the outcome of disease dynamics (5, 3, 2). For example, recent experimental studies indicate that the dramatic size of the 2013-2016 Ebola epidemic can at least be partially explained by the acquisition of genetic mutations that increased transmissibility to humans (6, 7).

Stochastic models of epidemiology can help to understand the emergence of evolving pathogen populations (5, 8–12). Available models, however, often make the unrealistic assumption that the pathogen is spreading in a well-mixed and homogeneous host population where all hosts are equally susceptible. Here we develop a general multi-host model to analyze the probability of evolutionary emergence of a pathogen in heterogeneous host populations where only some hosts are resistant to the pathogen. We demonstrate that realistic increases in the diversity of host resistance alleles strongly reduce the probability of evolutionary emergence of novel pathogens, hence suggesting new strategies to manage the emergence of diseases. Crucially, we developed a new experimental system using bacteria with distinct CRISPR (Clustered Regularly Interspaced Short Palindromic Repeat) immunity and their lytic viruses (bacteriophages) (13–16) to explore the effect of host population heterogeneity on the emergence and evolution of pathogens. The experimental validation of our theoretical predictions with this microbial system confirms the ability of our mathematical model to capture the complexity of the interplay between the epidemiology and evolution of emerging pathogens.

## Results

### Theory: predicting the probability of pathogen emergence

In order to predict how the composition of host populations impacts the probability of pathogen emergence, we developed a branching process model (5, 8–12). In this model, we assume that the host population contains a fraction (1 − *f_R_*)of individuals that are fully susceptible to the pathogen while the remaining fraction *f_R_* of the population is resistant and composed of a mixture of *n* host types in equal frequencies, each of which has a different resistance allele. The efficacy of resistance is governed by the parameter *ρ*. When *ρ* = 1, resistance is perfect and hosts can only be infected by pathogens that carry an escape mutation (one of which is available for each host resistance allele). When *ρ* < 1 host resistance is imperfect and (1 – *ρ*) host resistance is imperfect and measures the probability that a pathogen can cause an infection without the required escape mutation. Therefore, a pathogen with *i* escape mutations (*i* between 0 and *n*) can infect a fraction 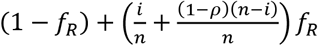 of the total host population.

We further assume that a host infected with a pathogen that does not carry escape mutations transmits at rate *b* and dies at rate *d*. Host resistance prevents infection without affecting *b* or *d*. Whereas escape mutations allow the pathogen to infect a larger fraction of the host population, they also carry a fitness cost *c* which causes pathogens with *i* escape mutations to reproduce at rate *b_i_* = *b*(1 – c)^*i*^. The probability of acquiring an escape mutation is a function of *n*, the number of resistance alleles in the population, as well as *i* the number of escape mutations already encoded by the pathogen. The probability that a pathogen with *i* escape mutations acquires an additional one equals *u_i,n_* = *μ*(*n* – *i*)/(*N* –*i*), where *μ* is the pathogen mutation rate per base pair, and *N* is its genome size. This simplifies to *u_i,n_* ≈ *μ*(*n* – *i*)/*N* when the pathogen genome is considerably larger than the number of escape mutations (i.e. *N* ≫ *i*). For the sake of simplicity, escape mutations cannot revert to the ancestral types. To approximate the effect of spatial structure we assume that when a parasite is released from an infected host it will land with probability *ϕ* on the same type of host (i.e. a susceptible host when *ρ* = 1), and with probability (1 − *ϕ*) on a random host from the rest of the population.

The expected number of secondary infections caused by a pathogen with *i* escape mutations in an uninfected host population is given by its basic reproduction ratio:

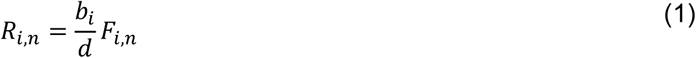

where *F_i,n_* = (*ϕ* + (1 − *ϕ*)( *f_R_* (*i* + (1−*ρ*)(*n*−*i*)) /*n* +(1 − *f_R_*))) is the effective fraction of hosts that can be infected by the focal pathogen. A pathogen with *n* escape mutations has a basic reproduction ratio equal to *R_n,n_* = *R*_0_ (1 − *c*)^*n*^, where *R*_0_ = *b*/*d* refers to the basic reproduction ratio of the unmutated pathogen in a fully susceptible host population.

The key question we wished to address with this model was how the composition and structure of the host population determines the ultimate fate of a pathogen (i.e. extinction versus emergence). We detail in the **Supplementary Information** the calculation of the probability of *emergence, P_i,n_*, which is the probability that an inoculum of *v*_0_ pathogens with *i* escape mutations does not go extinct when introduced in a host population with *n* different resistance alleles. To understand the role of pathogen evolution in this process, we also derive the probability of *evolutionary emergence*, which quantifies the importance of escape mutations to pathogen emergence.

### Evolutionary emergence is maximized for intermediate proportions of resistant hosts

To understand how the composition of the host population determines the probability that a pathogen can emerge, we consider the simple scenario where the population consists exclusively of sensitive hosts and one type of resistant hosts with the same resistance allele (i.e. *n* = 1). In this scenario, the probability of emergence of a phage with no escape mutation is (see **Supplementary Information**):

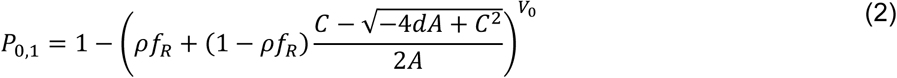

With *A* = *b* (1 − *μ*/*N*)( 1 − *ρf_R_* ( 1 – ϕ)), *B* = *bμ*/*N* and *C*= *A* + *B* (1 –1s/(*R*_0_(1 – *c*))) + *d*.

In the absence of mutation, the probability of pathogen emergence is thus:

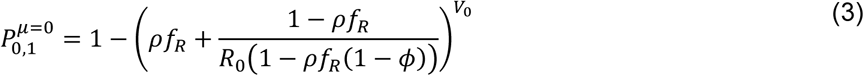

Varying the fraction *f_R_* of resistant hosts shows that the probability of emergence decreases linearly with the fraction of resistant hosts in well-mixed populations (when *ϕ* = 0 and *V_0_* = 1, **figure 1**). As expected, the probability of emergence decreases with *ρ*, the efficacy of host resistance, and increases with *V_0_*, the size of the pathogen inoculum (**figure 1**). Population structure has also a positive effect on pathogen emergence, since it helps the virus to persist in the susceptible subpopulation. Interestingly, there is a threshold value *f_T_* for the fraction of resistance where the probability of emergence vanishes (**figure 1**):

**Figure 1:**
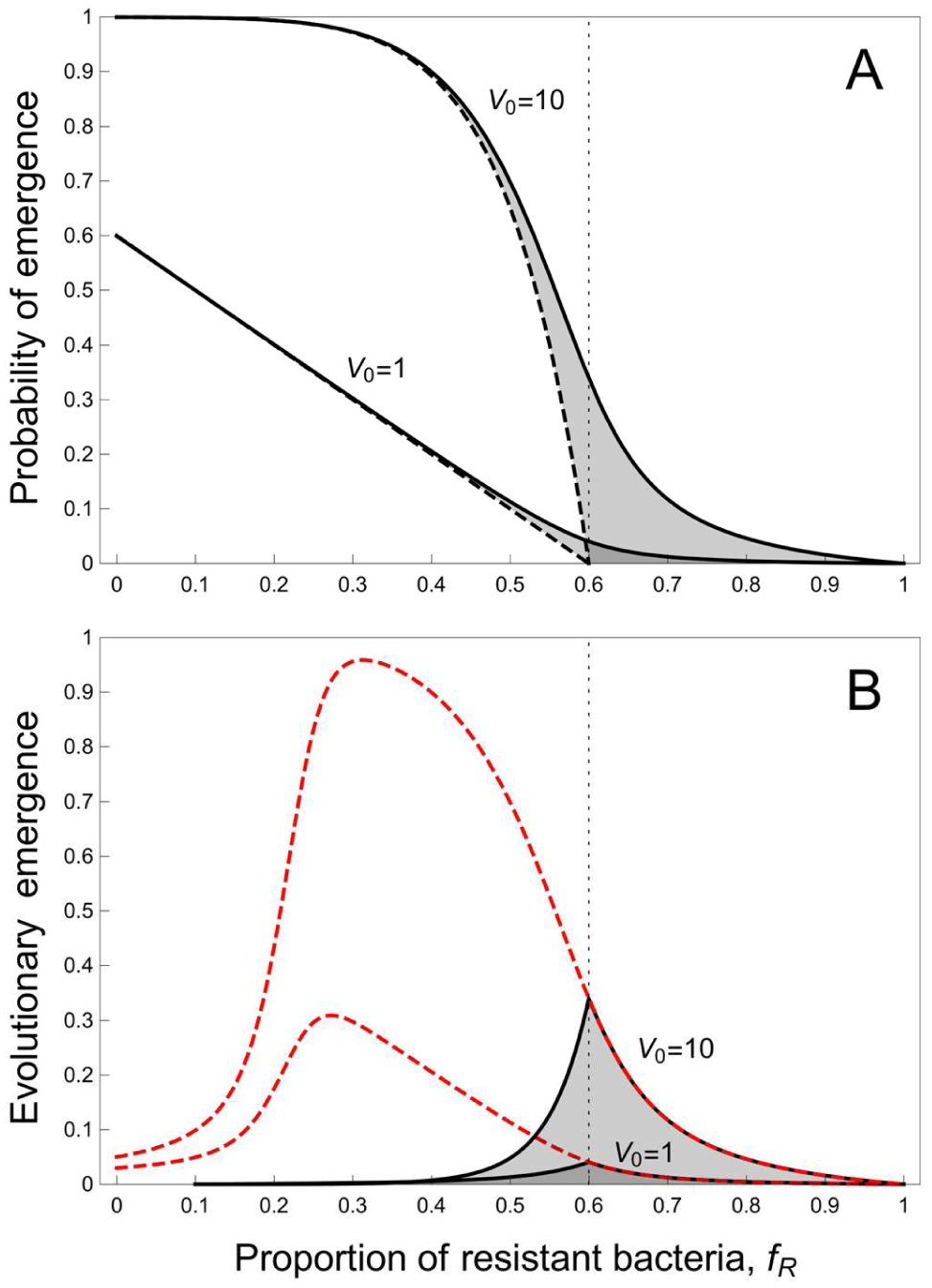
Effect of the proportion of resistant hosts (*f_R_*) on pathogen emergence when there is single type of resistant host (*n* =1) and for two values of the phage inoculum size (*V*_0_ = 1 and 10). (A) Probability of pathogen emergence without (*u*_0,1_ = *μ*/*N* = 0, dashed curve) or with (*u*_0,1_ = *μ*/*N* = 0.01, full curve) mutations. The shaded area illustrates to the fraction of pathogen emergence due to pathogen adaptation. The threshold value *f_T_*, of the fraction of resistant hosts that prevents pathogen emergence in the absence of pathogen adaptation, is indicated with a vertical dashed line. (B) Evolutionary emergence of pathogens (the shaded area in A) is maximized for an intermediate value of the fraction of resistant hosts. The dashed red curve represents the theoretical prediction when we track the change the frequency of escape mutations after the emergence (see **Supplementary Information**). Other parameter values: *b* = 2.5, *d* = 1, *ϕ* = 0, *ρ* = 1, *c* = 0.2, *T* = 24.

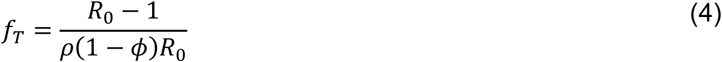

Next, we wanted to understand how pathogen evolution can help pathogens to emerge. As expected, pathogen mutation generally increases the probability of emergence (unless there is a significant fitness cost associated with escape mutations) because those mutations allow escape from host resistance. The gray area in **figure 1A** measures the amount of evolutionary emergence. Interestingly, evolutionary emergence is maximized for intermediate values of the frequency of resistance (usually when *f_R_* = *f_T_*, see **figure 1B**). Indeed, when the frequency of resistance is low there is no selection for escape mutations and those mutations get rapidly lost if they are associated with fitness costs. In contrast, when the frequency of resistance is high, the chains of transmission driven by the wild type strain are very short and there is a lower rate of appearance of escape mutations. Hence, in spite of the strong selection for escape mutations, the rate of evolutionary emergence is reduced. Intermediate resistance frequency therefore maximizes evolutionary emergence because this is where both the influx and the selection for escape mutations are high. When the efficacy of host resistance is low and when the host population is spatially structured, the role of pathogen evolution is smaller, simply because the probability of emergence without evolution is higher under these conditions (see **figures S1 and S2**).

So far, the model considered only the role of pathogen mutation during the very early stages of the epidemic, where stochastic effects play an important role. However, whenever we observe the emergence of a novel pathogen, these initial stages will usually have already passed, and pathogen evolution may therefore deviate from the patterns that are predicted by the above model. For example, even when escape mutations are not necessary for emergence, those mutations may increase in frequency after the onset of an epidemic. This is particularly true when the proportion of resistant hosts is large relative to the cost of mutation. After emergence, the size of the pathogen population increases rapidly and one can start to neglect the effect of demographic stochasticity on the change in escape mutation frequencies. To understand how pathogens evolve over longer timescales during an epidemic (i.e. beyond the initial emergence) we combined our stochastic description of pathogen emergence with a deterministic model for the change in the frequency of escape mutations (see **Supplementary Information**). This model predicts that the probability of observing an escape mutation evolving in a pathogen population is maximized for a low frequency of host resistance (red dashed curve in **figure 1B** and in **figures S1, S2** and **S3**).

### Diversity of host resistance decreases pathogen emergence

Natural host populations usually consist of multiple host genotypes, each carrying their own resistance allele. To understand how this will impact the predicted patterns of pathogen emergence, we analyzed the situation where the resistant host population consists of *n* types, at equal frequencies and each carrying a unique resistance allele. In the absence of pathogen mutations, emergence is governed by (3) and does not depend on host diversity because all hosts are equally resistant to a pathogen with no escape mutations. Each extra escape mutation, however, allows the pathogen to infect a fraction 1/*n* of the resistant host population, and the pathogen needs *n* escape mutations to exploit the whole host population. Yet, the probability *u_i,n_* to acquire an escape mutation increases with *n* because there are more genetic loci involved in the interaction with the host. In other words, both the number of mutations required to reach the top of the fitness landscape and the rate of acquisition of escape mutations increase with the diversity of host resistance. Because these two processes have opposite effects on the rate of pathogen adaptation, it is not immediately obvious how host diversity affects pathogen emergence.

We derive numerically the probability of pathogen emergence under a broad range of scenarios (see **Supplementary Information**). In **figure 2** we show the joint effects of the frequency of resistance and the diversity of resistance. Increasing host diversity always decreases the probability of emergence of the pathogen. We also find a very strong interaction between host diversity and spatial structure. Because spatial structure induces a clustering of the different types of resistant hosts it reduces the effective diversity of host resistance and promotes pathogen emergence (**figure S4**).

**Figure 2:**
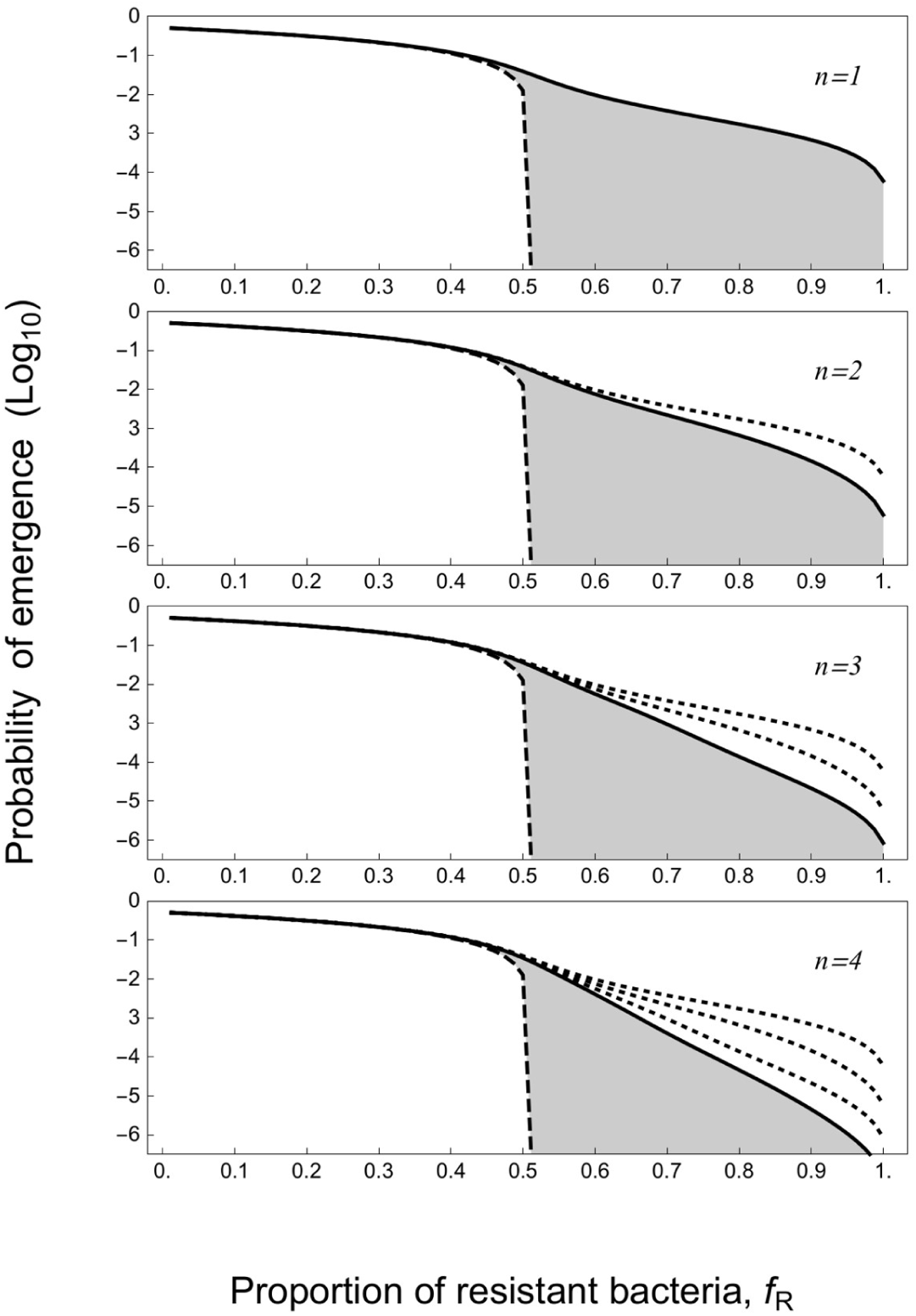
Effect of the fraction of resistant hosts and of host resistance diversity (*n*) on pathogen emergence. Each panel presents the probability of pathogen emergence without (*u*_0,*n*_ = *μ*/*N* = 0, dashed curve) or with (*u*_0,*n*_ = *μ*/*N=* 5 ∗ 10^−3^, full curve) mutations (note the logarithmic scale) for different values of host diversity (*n* = 1, 2, 3, 4). The shaded area illustrates the fraction of pathogen emergence due to pathogen adaptation. The dotted curves indicate the probability of emergence for lower levels of host diversity. Other parameter values: *b* = 2, *d* = 1, *ϕ* = 0, *ρ* = 1, *c* = 0.05.

The above analysis relies on (i) the assumption that the pathogen life-cycle can be approximated by the birth-death model, and (ii) the assumption that the frequency of host resistance does not vary through time. We relaxed both these assumptions with individual based simulations in the **Supplementary Information** and obtained very similar results (see **figures S5, S6** and **S7**).

### Experiments on phage emergence confirm theoretical predictions

Next we wanted to explore experimentally the validity of the above predictions. While this is challenging given the paucity of suitable empirical systems that are amenable to experimental manipulations, we explored whether this could be achieved by studying the evolutionary emergence of “escape” phages against bacteria with CRISPR-based resistance. The mechanism of CRISPR-based resistance relies on the addition of phage-derived sequences (known as “spacers”) in a CRISPR locus in the bacterial host genome (13). This empirical system allowed us to overcome three important technical challenges. First, the stochastic nature of extinction requires a large number of replicate populations to measure a probability of emergence, which is possible using bacteria and phages. Second, by mixing bacteria with different and unique CRISPR resistance alleles we could manipulate the fraction of resistant hosts and the diversity in resistance alleles without affecting other traits of the host (17). Third, unlike most other empirical systems to study host-pathogen interactions, the genetic mechanism that enables the pathogen (phage) to adapt to the host (bacterium) resistance through the acquisition of escape mutations is well understood for CRISPR-phage interactions, where lytic phages “escape” CRISPR resistance through mutation of the target sequence (the “protospacer”) on the phage genome (13, 15, 18, 19).

In order to validate the model using this empirical system, we used 8 CRISPR-resistant clones (also referred as <u>B</u>acteriophage Insensitive <u>M</u>utants, or “BIM”) of the Gram-negative *Pseudomonas aeruginosa* strain UCBPP-PA14, each of which carried one single spacer targeting the virulent phage DMS3vir. For each of these 8 CRISPR-resistant clones, the rate at which the phage acquires escape mutations was found to be approximately equal to 2.8*10^−7^ mutations/locus/replication, as determined using Luria-Delbruck experiments (see **Supplementary Information** and **figure S8**). Using these CRISPR-resistant clones, we first tested the theoretical prediction that the probability of emergence increases with the size of the virus inoculum (*V*_0_). To this end, 96 replicate populations, each composed of an equal mix of sensitive bacteria and a CRISPR-resistant clone, were each infected with on average 0.3, 3, 30, 300 or 3000 phages. After 24 hours, we measured the fraction of phage-infected bacterial populations in which emergence had occurred. Consistent with the model predictions, we observed that the higher the phage inoculum size, the higher the probability of emergence (**figure 3**, dashed line). In addition, we measured the fraction of populations where the phages had evolved to escape CRISPR resistance. Again, in accordance with the theory, we found that larger phage inoculi were associated with an increased evolution of phage escape mutations (**figure 3**, full line, Kendall, z = 3.416, tau = 0.784, p < 0.001). Furthermore, we obtained very similar results using a different empirical system consisting of the lytic phage 2972 and its Gram-positive bacterial host *Streptococcus thermophilus* DGCC7710. In this experiment, 96 populations composed of sensitive and a CRISPR-resistant host were infected with either 2, 20 or 200 phages. As above, we found that a higher phage inoculum led to both a higher probability of emergence and a higher probability of evolutionary emergence (**figure S9**).

**Figure 3:**
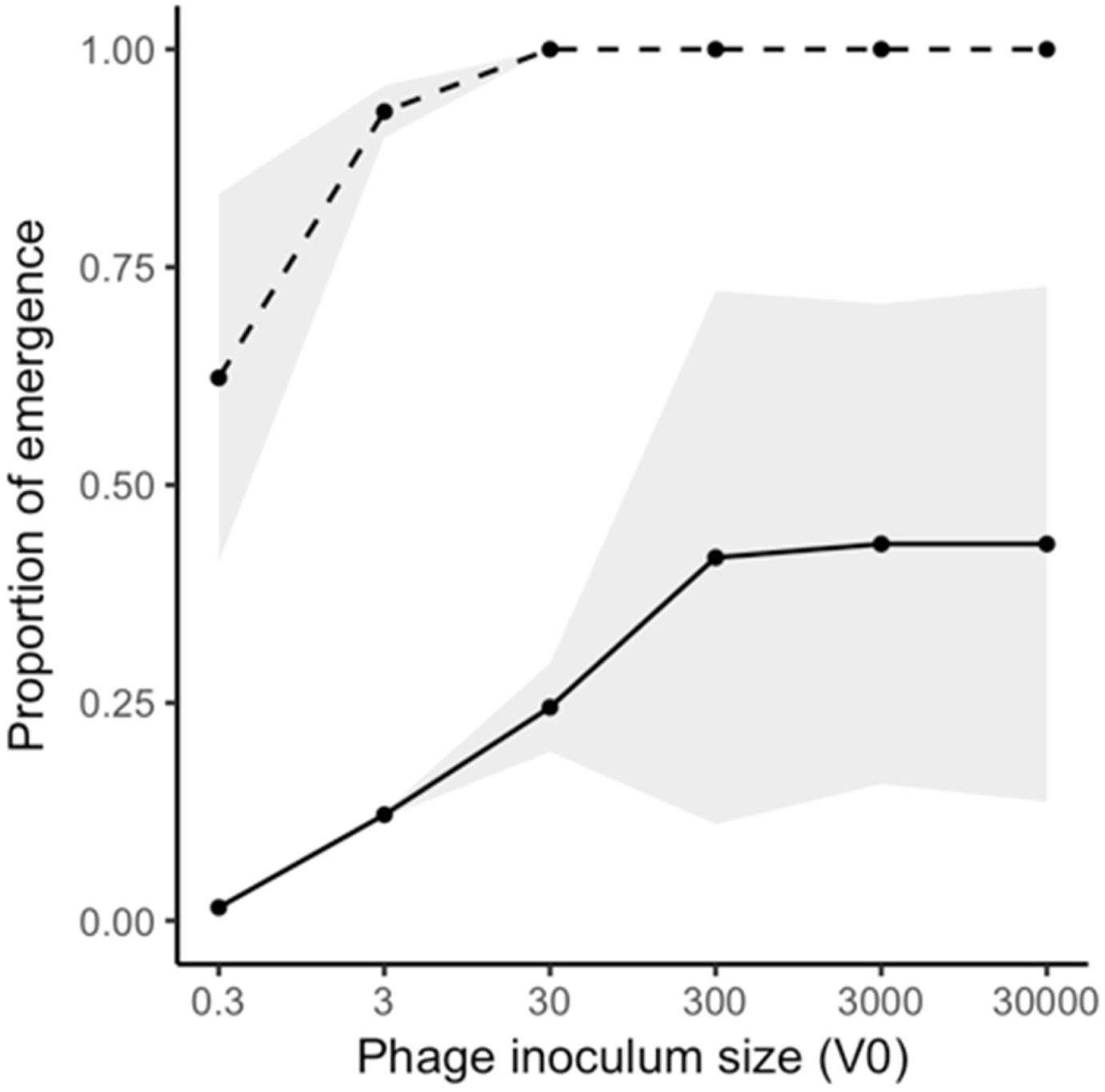
Probability of evolutionary emergence increases with the size of phage inoculum (*V*_0_). The proportion of replicate populations in which emergence (i.e. where the amplification of the phages is detected, dashed line) or evolutionary emergence (i.e. where the amplification of an escape phage is detected, solid line) was observed following infection with *V*_0_ ≈0.3, 3 30, 300 or 3000 unevolved phages in 96 independent replicate populations, each consisting of 50% sensitive bacteria and 50% BIMs ( *f_R_*= 0.5). Shaded areas represent 95% confidence intervals of the mean of two experiments.

Next, we tested the theoretical prediction that the probability of pathogen evolutionary emergence is highest in populations with an intermediate fraction of resistant hosts (**figure 4**). To this end, we generated populations composed of sensitive bacteria and a variable proportion of CRISPR-resistant bacteria, ranging from 0% to 100% in 10% increments. These populations were subsequently infected with *V*_0_ = 300 phages and the fraction of emergence and evolutionary emergence were measured. As expected, pathogen/phage emergence dropped when the proportion of host/bacteria resistance reached a certain threshold level (**figure S10**). Interestingly, examination of phage evolution among emerging phage populations also confirmed that the probability of observing escape mutations is maximized for intermediate proportions of host resistance (**figure 4**). Again, we obtained very consistent results with phage 2972 and *S. thermophilus* (**figure S9**).

**Figure 4:**
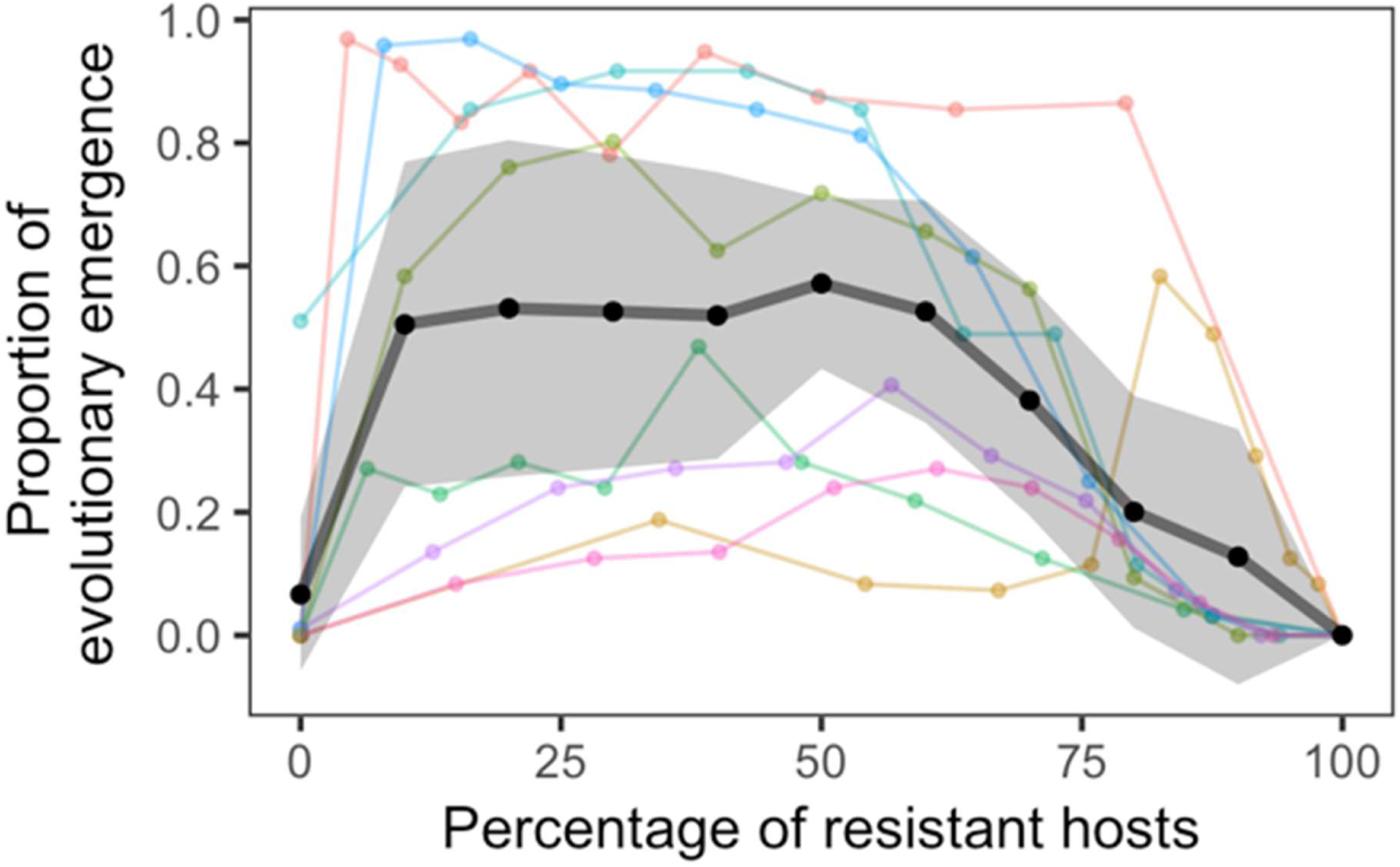
Intermediate proportion of resistant hosts maximises the probability of evolutionary emergence. Proportion of replicate populations with evolutionary emergence (i.e. where the amplification of an escape phage is detected) for increasing values of the proportion of resistant bacteria (*f_R_*). The different colors correspond to replicate experiments performed using 8 different BIMs (see **table S2** and **figure S8**). For each treatment, each of the 96 replicate host populations was inoculated with an initial quantity of *V*_0_ ≈300 unevolved phages. Black lines indicate the mean across the 8 BIMs; grey shaded areas represent 95% confidence intervals of the mean.

We noticed substantial variation among CRISPR-resistant hosts in the observed frequencies of escape phage evolution (**figure 4**). Variations in phage mutation rates are unlikely to explain this variability because, as pointed above, we failed to detect significant variations in the rate of escape mutations to the different CRISPR-resistant hosts (see **figure S8**). Variations in the fitness cost associated with these mutations could, however, explain the observed variations in the final frequency of escape mutations (see **figure S3**).

Finally, we experimentally explored the effect of resistance allele diversity on evolutionary emergence for a fixed proportion of host resistance (*f*_R_ = 0.5). To this end, we generated bacterial populations that were composed of sensitive bacteria and an equal mix of 1, 2, 4 or 8 CRISPR-resistant clones. In this case, as expected, an inoculum size of 300 phages always led to pathogen emergence, but increasing host diversity had a strong negative effect on the ability of the phage to evolve to escape host resistance (**figure 5**). We also found higher probabilities of observing multiple escape mutations in the low diversity treatment (Kendall, z = - 4.8771, Tau = −0.3259, p = 1.07*10^−6^), which also supports the prediction that host diversity hampers the evolution of the phage population.

**Figure 5:**
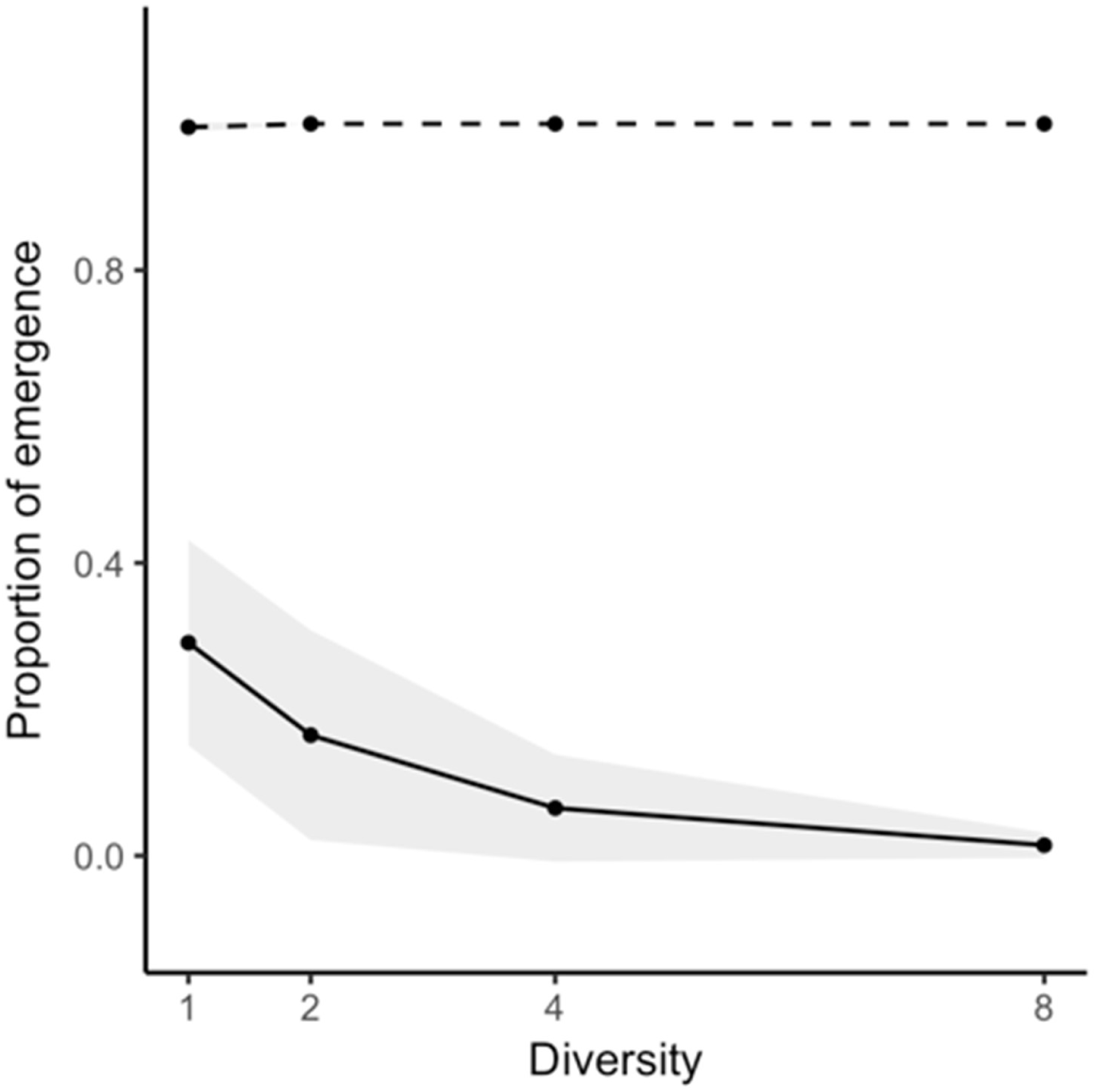
Increasing the diversity of host resistance decreases the probability of evolutionary emergence. Proportion of replicate populations with phage emergence (i.e. where the amplification of the phages is detected, dashed line) or evolutionary emergence (i.e. where the amplification of an escape phage is detected, solid line) for increasing values of the diversity of host resistance (*n* =1, 2, 4 or 8 BIMs) when the proportion of host resistance is *f_R_* = 0.5. For each treatment, each of the 96 replicate host populations was inoculated with an initial quantity of *V*_0_ ≈300 unevolved phages. Shaded areas represent 95% confidence intervals of the mean of two experiments.

## Discussion

The emergence and re-emergence of pathogens has far-reaching negative impacts on wildlife, agriculture and public health. Unfortunately, pathogen emergence events are notoriously difficult to predict and we need good biological models to experimentally explore the interplay between epidemiology and evolution taking place at the early stages of an epidemic. Here we used a combination of diverse theoretical and experimental analyses to examine how the composition of a host population impacts the probability of pathogen emergence and evolution. Our experimental validation of these predictions using microbial populations demonstrates the predictive power of this theoretical framework and its relevance for the control of emerging infectious diseases.

Our framework provides important insights regarding emergence and re-emergence in both the presence and absence of pathogen evolution. For instance, this model captures how the composition and diversity of the host population impacts the emergence of a non-evolving pathogen. In this context, a larger proportion of resistant hosts decreases pathogen emergence, but this effect is weaker in spatially structured populations where transmission is more likely to occur between the same host types, which allows for pathogen persistence in sensitive subpopulations. This effect is akin to the effect of the spatial distribution of suitable habitats on extinction thresholds (20–23) and consistent with earlier work which shows that host composition and spatial structure impact the growth rate of bacteriophage ϕ6 (24). In the context of an evolving pathogen, our theory helps to explain the general observation that evolutionary emergence and the spread of escape mutations is maximal for an intermediate proportion of resistant hosts in the population (25). Specifically, this is because increasing host resistance in the population has two opposite effects: (i) the influx of new mutations decreases because the ancestral pathogen cannot replicate on resistant hosts, (ii) selection for escape mutations increases.

Second, our model predicts that diversity in host resistance alleles decreases the probability of evolutionary emergence. Even though larger host diversity increases the number of adaptive mutations for the pathogen (i.e. a larger number of targets of selection), each mutation is associated with a smaller fitness advantage (i.e. a smaller increase in the fraction of the host population that can be infected). The theory presented here therefore helps to explain previous empirical data on the impact of host CRISPR diversity on the evolution of escape phages (17). The link between host biodiversity and infectious diseases has attracted substantial attention recently (26–37). Several studies support the “dilution effect” hypothesis which postulates that host diversity limits disease spread (34–36). For example, host diversity may limit the spread of a pathogen by increasing the fraction of bad quality hosts in the population (36). Indeed, increasing the fraction of resistant hosts (but not the diversity of resistance alleles) decreases the basic reproduction ratio of the wild type pathogen (38, 39). In addition, host diversity per se may also limit disease spread and several studies have shown the negative effect of host diversity on the deterministic growth rate of the pathogen under specific patterns of host-parasite specificity (29, 40, 41).

Notwithstanding these important insights, what sets our theoretical model apart is its ability to predict the factors that impact the initial pathogen emergence, rather than the downstream spread of a pathogen once it has already emerged. Studying this requires stochastic models, which are critical to model the probability of rare events, for example pathogen spillover across species, including at the Human-Animal interface (42, 43, 3, 44), the emergence of drug resistance (45, 46), the evolution of vaccine resistance (47) and the reversion of live vaccines (48–52). In all these public-health issues, predicting pathogen emergence requires models accounting for the stochastic nature of epidemiological and evolutionary dynamics. Our work provides a theoretical framework to study these different issues and can thus be used to evaluate the ability of different control strategies to limit pathogen adaptation and emergence.

## Acknowledgments

The authors thank Jack Common for providing the different *P*. *aeruginosa* BIMs. HC acknowledges funding from the EMBO (Short Term Fellowship program). Sean Meaden was supported by funding from the BBSRC (BB/N017412/1), which was awarded to ERW. ERW further acknowledges the Natural Environment Research Council (NE/M018350/1), the Wellcome Trust (109776/Z/15/Z) and the European Research Council (ERC-STG-2016-714478 -EVOIMMECH) for funding. SVH acknowledges funding from the People Programme (Marie Curie Actions; https://ec.europa.eu/research/mariecurieactions/) of the European Union’s Horizon 2020 (REA grant agreement no. 660039) and from the BBSRC (BB/R010781/1). LMW and SM acknowledge funding from the Natural Sciences and Engineering Research Council of Canada. SM holds the Tier 1 Canada Research Chair in Bacteriophages. SG acknowledges support from the CNRS (PEPS MPI grant) and the Leverhulme Trust (Visiting Professorships grant).

